# P-NADs: *P*UX-based *NA*nobody Degraders for Ubiquitin-Independent Degradation of Target Proteins

**DOI:** 10.1101/2023.12.20.572486

**Authors:** Jun Wang, Georgy Chistov, Junrui Zhang, Brandon Huntington, Israa Salem, Anandsukeerthi Sandholu, Stefan T. Arold

## Abstract

Targeted protein degradation (TPD) allows cells to maintain a functional proteome and to rapidly adapt to changing conditions. Methods that repurpose TPD for the deactivation of specific proteins have demonstrated significant potential in therapeutic and research applications. Most of these methods are based on proteolysis targeting chimaera (PROTAC) which link the protein target to an E3 ubiquitin ligase binding moiety, resulting in the ubiquitin-based degradation of the target protein. In this study, we introduce a method for ubiquitin-independent TPD based on nanobody-conjugated plant ubiquitin regulatory X domain-containing (PUX) adaptor proteins. We show that the *P*UX-based *NA*nobody *D*egraders (P-NADs) can unfold a target protein through the Arabidopsis and human orthologues of the CDC48 unfoldase without the need for ubiquitination or initiating motifs. Despite originating from plants, P-NAD plasmids can be transfected into a human cell line, where produced proteins use the endogenous CDC48 machinery for ubiquitin-independent TPD. Thus, P-NADs pave the road for ubiquitin-independent therapeutic TPD approaches. The P-NAD design combined with established *in vitro* and cellular assays make this system also a versatile platform for elucidating functional aspects of CDC48-based TPD in plants and animals.

## INTRODUCTION

Cells of all organisms are tasked with two critical functions. Firstly, they must eliminate dysfunctional proteins to maintain a functional proteome and ensure homeostasis. Secondly, they must swiftly adapt their proteome in response to changing internal or external needs and signals. Both these functions can be accomplished at the protein level through Targeted Protein Degradation (TPD). TPD holds significant potential as a research tool for decoding protein function, and as a therapeutic strategy for degrading proteins with pathogenic functions (Luh et al., 2020).

The concept of Proteolysis Targeting Chimeras (PROTACs) has emerged from the understanding that cells mark proteins for Targeted Protein Degradation (TPD) by ubiquitinating them. This process culminates in the transfer of ubiquitin to the target protein via an E3 ligase. PROTACs are adapters that connect target proteins to an E3 ligase, facilitating their poly-ubiquitination and subsequent degradation through the cellular proteasome-dependent pathway. PROTACs have two functional components: one that binds to the protein of interest, and another that attaches to specific E3 ligases (Sakamoto, 2001). Initial PROTACs were small chemical entities, limiting their application to E3 ligases and target proteins for which specific small-molecule inhibitors were available. More recently, PROTACs have been developed where the protein of interest is captured through a nanobody (a binding domain derived from single-chain antibodies) (Caussinus, 2011; Morimoto, 2015; Clift, 2017; Baudisch, 2018; Ibrahim, 2020). Given that nanobodies can be obtained against many proteins, these nanobody degraders enable the creation of PROTACs for proteins that could not be targeted by small molecules.

However, nanobody-based degraders still depend on the ubiquitination machinery of the target cell, introducing some limitations: (i) The target protein must have a well-positioned lysine residue on a surface site accessible to the E2 ubiquitin-conjugating enzyme (W. Zhang, 2022). (ii) Most organisms host a plethora of E3 ligases in their genome (> 600 in humans), each with highly different structures and features (Li, 2008). The expression of a specific E3 ligase can vary significantly across different cell types, necessitating the development of different PROTACs for different tissues or organisms. (iii) The nanobody PROTACs are also proteins, which means this system can lead to the ubiquitination and degradation of the PROTAC before it can bind to its target. (iv) Finally, ubiquitinated proteins run the risk of forming aggregates (Morimoto et al., 2015).

In this study, we introduce a novel type of degrader based on Plant Ubiquitin X Regulatory (UBX) domain-containing proteins (PUX) (Kaneko, 2021). UBX proteins function as adaptors to the segregase and unfoldase Cell Division Control Protein 48 (CDC48) (Schuberth, 2008). CDC48 is a hexameric AAA ATPase that is highly conserved, with almost 80% sequence identity across yeast, plant, and animal orthologues. CDC48 employs a variety of species-specific UBX proteins as adapters to extract and unfold target proteins, which are then degraded through the proteasome (Kloppsteck, 2012).

We demonstrate that degraders, constructed from a combination of nanobodies and PUX proteins from *Arabidopsis thaliana*, can promote CDC48-dependent protein degradation even in the absence of ubiquitination (**Figure 1A**). This addition to the TPD toolkit opens up new possibilities for ubiquitin-independent TPD in research or therapeutic applications.

**Figure 1.**
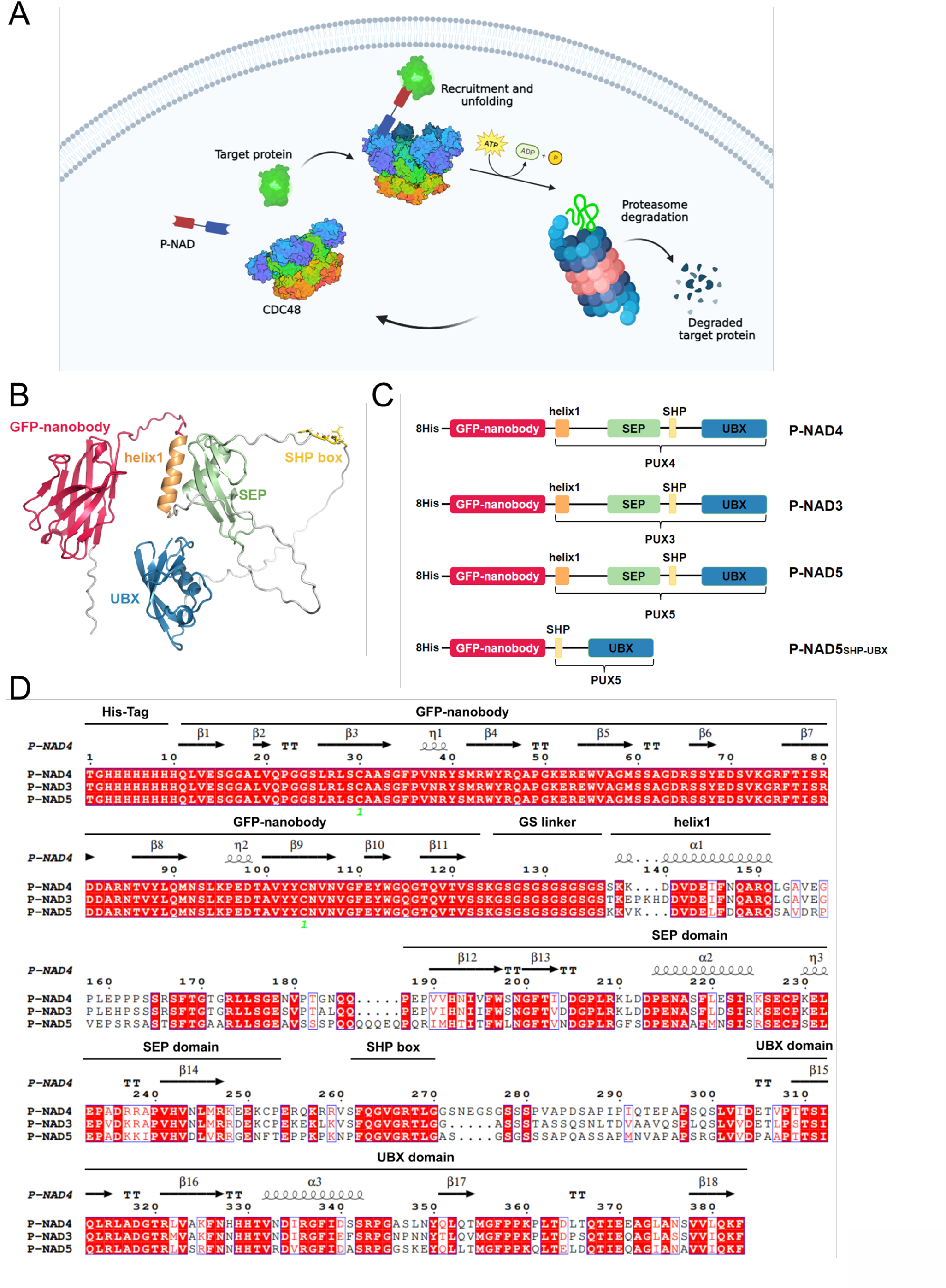
Targeted protein degradation by P-NADs. (**A**) Schematic diagram for P-NADs-mediated protein degradation. Figure was generated in BioRender.com. P-NADs bind their target proteins *via* the N-terminal nanobody, and recruit CDC48 through their C-terminal domains, forming a ternary complex. CDC48 then unfolds the target protein, facilitating proteasomal degradation. (**B**) AlphaFold structure prediction of P-NAD4. (**C**) Domain architecture of P-NADs designed in this study. (**D**) Multiple sequence alignment of P-NAD3, 4, and 5, showing the predicted secondary structure, sequence motifs and protein domains. The alignment was performed by ClustalW and visualised by ESPript3.

## RESULTS

### Design of Nanobody-UBX Degraders

For our TPD design, we selected the PUX CDC48 adaptors as the basis. For nomenclature, we use CDC48 to refer to all homologues in all organisms, CDC48A to refer to the primary orthologue in *A. thaliana*, and p97 for the human orthologue. Across organisms, most UBX proteins use their C-terminal UBX domain to bind to the N-terminal domain of CDC48, while their N-terminal region binds to target proteins (Buchberge, 2015). Many UBX proteins contain additional central domains of which the functions are not completely understood (Bashore, 2023; J. Zhang, 2022).

The use of PUX proteins as a basis for ubiquitin-independent TPDs was motivated by the following: (i) PUX proteins and their yeast and animal orthologues constitute the largest family of CDC48 cofactors (Arabidopsis alone contains 16 PUX proteins (Rancour, 2004), and hence the largest basis for TPD design templates; (ii) PUX proteins are singly-chain adapters, whereas the other prominent CDC48 adaptor is a more complex heterodimer formed by UFD1 and NPL4 (Bruderer, 2004); and (iii) PUX proteins have a modular “beads-on-string” architecture that is convenient for protein engineering. Based on our interest in designing a ubiquitination-independent TPD, we chose PUX3, PUX4 and PUX5 because of their dual, and hence more stable, linkage to the N-domain of CDC48A through a UBX domain and the ten-residue SHP box motif (Zhang, 2021)(**Figure 1B**).

Moreover, PUX3, 4 and 5 can be assigned to the same Group IV as the human UBX protein p37, based on analogies in their domain composition (Zhang, 2022). The p37:p97 pair has been shown to disassemble the PP1:SDS22:I3 complex in a ubiquitin-independent manner (Weith et al., 2018). In p37, this process required the presence of the SEP domain and preceding helix (named hereafter helix1; (Weith, 2018), and PUX3, 4, and 5 also contain homologues of this feature (Zhang, 2022). We therefore used the fragments including the helix1-SEP-SHP-UBX regions of PUX3, 4 and 5 as basis (**Figure 1C**). The sequence identity of these PUX proteins is approximately 60% (**Figure 1D**). To facilitate assessing unfolding *in vitro* and in cells, we chose a well-characterized nanobody targeting GFP (Kubala, 2010), which we N-terminally fused to the PUX bases using a twelve amino acid flexible linker (**Figure 1D**). We added an N-terminal hexahistidine tag to facilitate purification. The resulting *P*UX-based *NA*nobody *D*egraders (P-NADs, pronounced /’pi?.n∧ts/) P-NAD3, P-NAD4, and P-NAD5 (named according to their base being PUX3, 4, or 5) were 384 ± 4 residues long (**Figure 1C**,**D**). AlphaFold models and their predicted aligned errors plots (PAEs) were consistent with a “beads-on-string” model where all domains are flexibly linked, except for helix1 which was predicted to bind the SEP domain in all P-NADs (**Supplementary Fig 1A, B**) (Zhang, 2022). To probe the influence of the helix1-SHP fragment, we created an additional fragment for PUX5 where the helix and SEP domain were deleted (P-NAD5_SHP-UBX_, **Figure 1C**).

### Recombinant P-NADs bind to CDC48A, p97 and target

We recombinantly produced all four P-NADs through protein expression in *E. coli* followed by purification through nickel columns and size exclusion columns (**Methods**). Protein yields (5-15 mg per L of culture) and purity were satisfactory (**Supplementary Figure 2A**). Size exclusion elution volumes corroborated that P-NADs were mostly monomeric molecules (**Supplementary Figure 2B**). We used ITC to test the interaction between P-NAD4 and *Arabidopsis* CDC48A as well as its interaction with human p97. Despite its origin from plants, P-NAD4 bound to p97 and CDC48A-ND1 (a hexameric CDC48 construct containing the N and first ATPase domains) with similar dissociation constants, *K*_*d*_*s*, of 6.14 ± 0.8 μM and 3.13 ± 0.6 μM, respectively (**Fig 2A, Supplementary Figure 3A**,**B**). This compatibility suggested that P-NADs could function in both mammalian and plant systems. The affinities of other wild-type or engineered constructs towards CDC48A-ND1 were comparable (*K*_*d*_ of 5.6 ± 1.6 μM, 2.7 ± 0.9 μM and 12.3 ± 3.7 μM for Arabidopsis PUX3, PUX4 and P-NAD5_SHP-UBX_; **Fig 2A, Supplementary Figure 3C-E**). The GFP nanobody bound to monomeric ultrastable GFP [muGFP (Scott, 2018), hereafter termed GFP for simplicity] with comparable nanomolar affinity alone or as part of a P-NAD construct, suggesting that the nanobody is not occluded by the construct design (**Fig 2A, Supplementary Figure 3F, G**).

**Figure 2.**
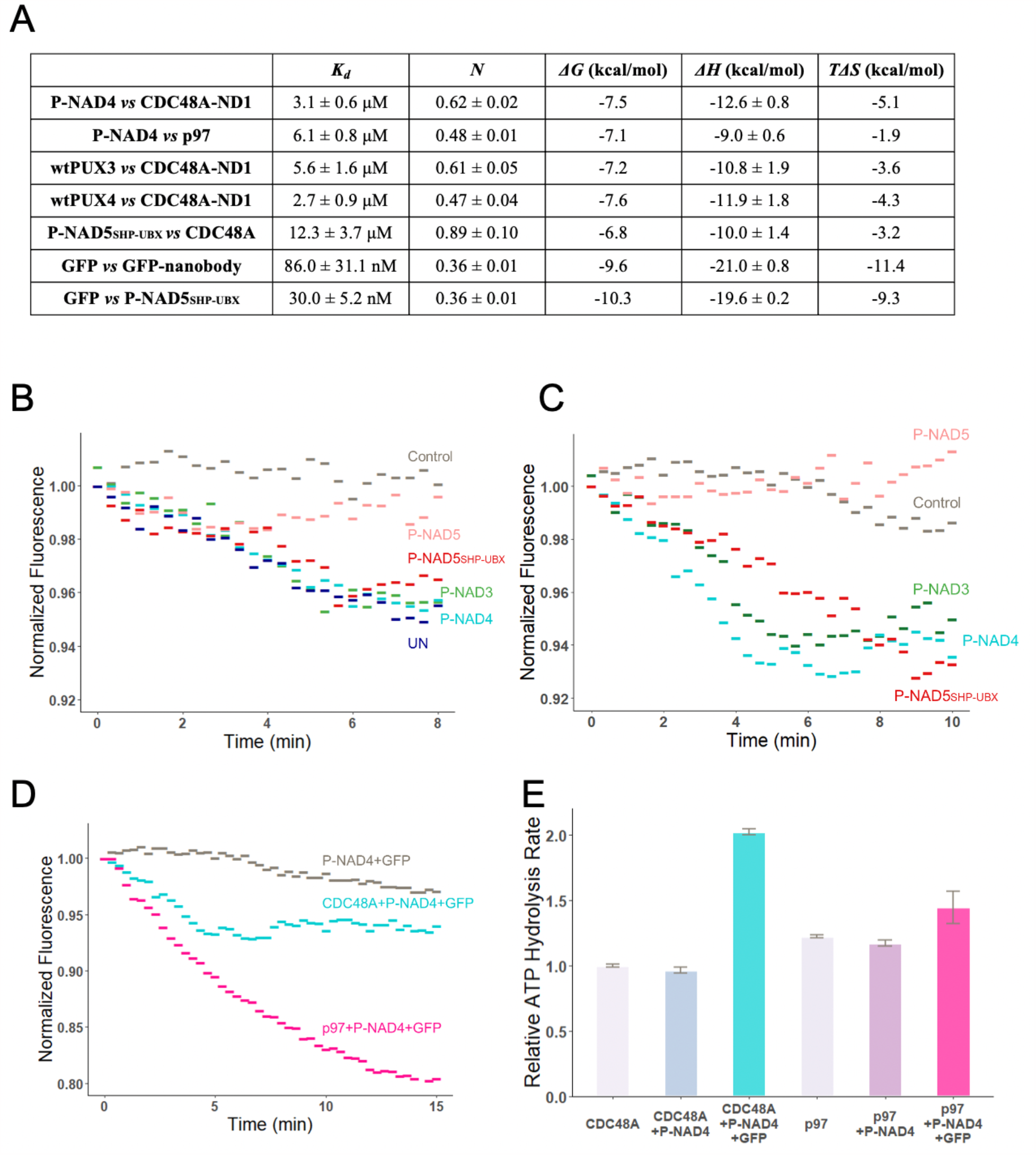
P-NADs bind GFP and CDC48A/p97, and induce GFP unfolding *in vitro*. (**A**) Table of ITC results obtained from the curve fit of data shown in **Supplementary Figure 3A-G**. (**B-C**) Fluorescence unfolding assay of GFP by P-NADs. GFP, P-NAD4, and CDC48A/p97 were incubated in unfolding assay buffer with ATP (see **Methods**). As a negative control, GFP was incubated with P-NAD4 without CDC48. (**B**) Unfolding of Ub-GFP by P-NADs. As a positive control, Ub-GFP was incubated with UFD1B-NPL4 and CDC48A. (**C**) Ubiquitin-independent unfolding of GFP by P-NADs and CDC48A. (**D**) Comparison of the unfolding efficacy of P-NAD4 with either CDC48A or p97. (**E**) ATP hydrolysis rates of CDC48A/p97 measured in the presence or absence of P-NAD4 and GFP. The rates were normalised to that of CDC48A alone in reaction buffer with ATP.

### P-NADs and CDC48/p97 unfold GFP *in vitro*

To monitor the capability of unfolding target proteins, we established an *in vitro* fluorescence unfolding assay (see **Methods**). First, we tested unfolding of a ubiquitinated substrate, which is considered the canonical type of substrates for CDC48. We established an *in vitro* polyubiquitination assay based on the use of the E3 ligase PRT1 from *Arabidopsis* (Stary, 2003), which is smaller and hence easier to obtain recombinantly and handle *in vitro* than the other canonical E3 ligases. We confirmed the production of polyubiquitinated GFP (Ub-GFP) by SDS-PAGE gel (**Supplementary Figure 3C, D**).

Incubation of Ub-GFP with CDC48A and P-NADs resulted in a decrease in GFP fluorescence over 5 minutes, until reaching a steady-state (**Figure 2B**). This steady-state fluorescence may correspond to the equilibrium between CDC48-promoted unfolding and spontaneous refolding of GFP. Indeed, GFP is known to have a strong capability to refold *in vitro* (Blythe, 2017; Bodnar, 2017). The unfolding rate and maximum varied between constructs, with P-NAD3 and P-NAD4 unfolding GFP markedly faster and more efficiently than P-NAD5. Interestingly, deleting the helix1-SEP module increased the unfolding activity of the remaining P-NAD5_SHP-UBX_ fragment, suggesting that the low unfolding efficacy of P-NAD5 is due to motifs within the helix1-SEP, at least *in vitro* (**Figure 2B**). As positive control, we used the *Arabidopsis* UFD1:NPL4 heterocomplex, which is a distinct ubiquitin-dependent adaptor for CDC48 (Ye, 2003). Ub-GFP degradation mediated by UFD1:NPL4 reached similar efficiencies than P-NAD3 and 4 (**Figure 2B**). GFP incubated with P-NAD4 without CDC48A was used as a control to monitor the fluorescence decay without active degradation.

We next probed the capability of the different adaptors to degrade GFP without ubiquitination. The presence of P-NAD3 or P-NAD4 promoted a marked fluorescent decrease of GFP, with similar or better efficacy than measured for Ub-GFP (**Figure 2C**). P-NAD5_SHP-UBX_, but not P-NAD5, showed significant degradation of GFP, in line with the activity of these constructs on Ub-GFP. We chose the best performing degrader, P-NAD4, to compare its performance between CDC48A and p97. Unexpectedly, the p97 unfoldase was markedly better in degrading GPF in presence of P-NAD4 than CDC48A (**Figure 2D**). This observation was surprising, because CDC48A had a significantly higher ATP hydrolysis rate in presence of P-NAD4 (**Figure 2E**).

We concluded that P-NAD3 and P-NAD4 can degrade a very stably folded target protein *in vitro*, even in the absence of ubiquitination and initiating motifs, and that p97 performed better than CDC48, at least in complex with P-NAD4. The performance of P-NAD5 was poor but could be enhanced by the removal of the helix1-SHP fragment.

### P-NADs promote GFP degradation *in vivo*

Finally, we investigated the function of the best-performing P-NAD in a living human cell. For this, we transfected both P-NAD4 and GFP into HeLa cells (**Figure 3**). Fluorescence imaging revealed that the presence of P-NAD4 led to a significant decrease in fluorescence intensity (**Figure 3A-C**). This observation was made in comparison to a control group that was only transfected with GFP, indicating that the introduction of P-NAD4 decreased the abundance of folded GFP. Western blotting validated the results by showing that the level of GFP decreased by 45% when P-NAD4 was present (**Figure 3D, E**). Importantly, the transfected GFP plasmid does not encode for a degron sequence (short linear motifs that mediate protein ubiquitination by binding to E3 ubiquitin ligases, thus triggering TPD by the ubiquitin-proteasome system (Sakamoto, 2001). Hence, the GFP produced in the HeLa cells is not ubiquitinated, as confirmed by a single band on the GFP western blot (compare **Supplementary Figures 2C** with **2E**). Consequently, P-NADs are also capable of promoting ubiquitin-independent TPD in HeLa cells.

**Figure 3.**
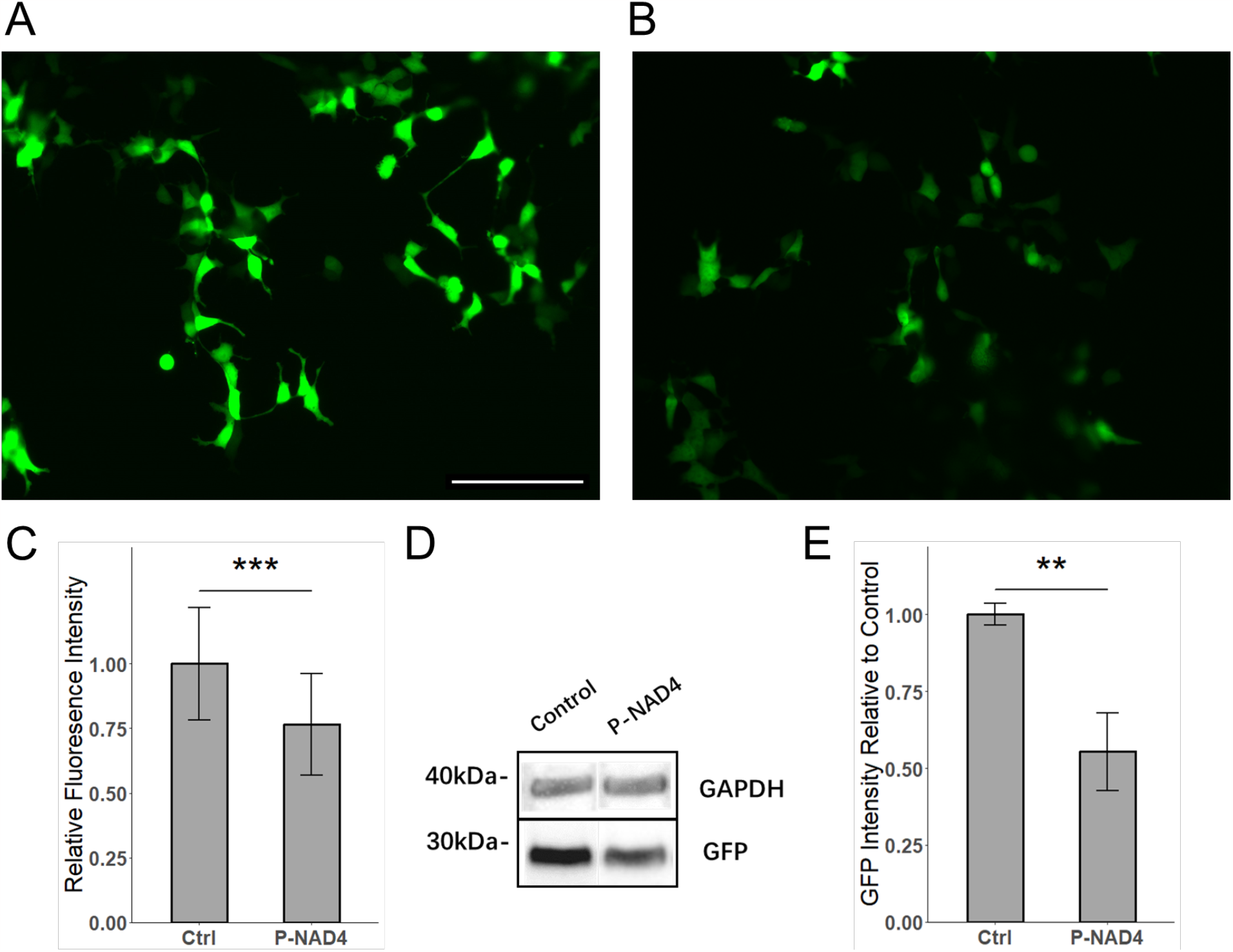
P-NAD4 induces GFP degradation in cells. Fluorescence microscopy images of HeLa cells transfected with GFP alone (**A**) or GFP and P-NAD4 (**B**). Scale bar: 150 μm. (**C**) Quantification of the GFP fluorescence intensity in the microscopy images by ImageJ software. The data are presented as the mean ± standard deviation of three independent experiments. Asterisks indicate significant differences from the control group (GFP alone) by Student’s t-test (*p < 0.05, **p < 0.01, ***p < 0.001). (**D**) Western blot analysis of the total cell lysates, probed with an antibody against GFP. The blot shows the protein level of GFP and the loading control (GAPDH). The molecular weight markers are indicated on the left. (**E**) Quantification of the western blot results by ImageJ software. The data are presented as the mean ± standard deviation of three independent experiments. Asterisks indicate significant differences from the control group (GFP alone) by Student’s t-test (*p < 0.05, **p < 0.01, ***p < 0.001).

Together, these results demonstrated that P-NAD4 can use endogenous p97 to promote targeted TPD of GFP in a human cell line. The actual efficacy of P-NAD4 degradation may be higher, because not all cells containing GFP are expected to be also co-transfected with P-NAD4.

## DISCUSSION

Various well-performing PROTAC variants have been reported, including ones that use nanobodies as target recognition modules (Caussinus, 2011; Morimoto, 2015; Clift, 2017; Baudisch, 2018; Ibrahim, 2020). However, these systems depend on ubiquitination of the substrate and hence on the cell- and organism-specific availability of E3 ligases. We designed P-NADs as a novel class of nanobody degraders that achieve TPD by linking target proteins to CDC48 unfoldases without the need for ubiquitination. Although our P-NADs are based on *Arabidopsis* PUX proteins, P-NAD4 also functions with human p97 *in vitro* and in cells. Indeed, despite being encoded by species that separated ∼1 billion years ago, both CDC48 orthologues remain almost 80% identical in their sequence. Hence, P-NADs are expected to be able to use the endogenous CDC48 unfoldase machinery from a large variety of organisms. Being in addition independent of cell type–specific E3 ligases, P-NADs have the promise to be compatible with many organisms and tissues.

The unfolding efficacy in our *in vitro* assays was markedly lower than the 45% reduction observed in transfected HeLa cells. We propose that this difference is due to the lack of proteasomes in our *in vitro* assay, which allowed the GFP to refold. Although the unfolding ratio reached *in vitro* was sufficient for a comparative study of degraders, the introduction of proteasomes, or the use of other means of blocking refolding may significantly increase the unfolding signal.

The integration of *in vitro* and cellular assays enables the utilisation of P-NADs to explore the not fully understood functionalities of the various domains and motifs of UBX domain adaptor proteins (Zhang, 2022). For instance, our findings indicate that P-NADs can facilitate the unfolding of a non-ubiquitinated GFP by CDC48. As of now, the disassembly of the PP1:SDS22:I3 complex by p97:p37 represents the sole known instance of CDC48-mediated ubiquitin-independent unfolding. In this mechanism, p97 identifies an internal recognition signal (IRS) in I3 and translocates it through the pore in a loop configuration rather than a single strand (van den Boom, 2021). Contrary to I3, which is an unstructured protein, our muGFP lacks an IRS and is ultra-stably folded (Scott, 2018). This result demonstrates that CDC48 can directly unfold well-structured proteins without requiring ubiquitin modification or initiating motifs, and that simply linking the substrate protein and CDC48 is sufficient to trigger unfolding. Future studies employing nanobodies aimed at different targets will yield further insights into the target features that are compatible with this form of TPD. Additionally, we observed that the helix1-SEP domain modules were not required for ubiquitin-independent TPD by P-NAD5. In fact, their presence reduced TPD by this P-NAD, implying an (auto)inhibition through an unknown mechanism.

The stoichiometry *N* of approximately 0.5 in titrations between hexameric CDC48 (CDC48A-ND1 or p97) and PUX or P-NADs (**Figure 2A**) suggests that three molecules of P-NUD bind to one CDC48/p97 hexamer. This stoichiometry may indicate that one P-NAD molecule can make contacts with two CDC48 N domains simultaneously, through the UBX domain and SHP box motif(s). The only exception here is P-NAD5_SHP-UBX_, which binds CDC48A-ND1 in a 1:1 complex, suggesting that the binding stoichiometry is influenced by the helix1-SEP fragment. PUX3, 4 and 5 contain an additional putative SHP box N-terminal to the SEP domain (F^168^XGXGRXL in P-NAD4, see **Figure 1D**) which might contribute to binding (Zhang, 2022). However, more data is required to confirm the stoichiometry and its molecular basis.

The compatibility of P-NADs with CDC48 systems across different kingdoms enables their use in investigating the unique characteristics of various CDC48 orthologues. For instance, we noted that p97 and CDC48A bound P-NADs with comparable affinity; yet, p97 outperformed CDC48 in P-NAD4–mediated unfolding, although p97 hydrolysed less ATP in presence of P-NAD4. Other less understood attributes, such as the adaptor-to-CDC48 stoichiometry, the necessary features for unfolding stable 3D domains, or the synergistic or competitive interactions between different types of CDC48 adaptors can be investigated with this system. As a result, our P-NADs offers a valuable resource for advancing our mechanistic comprehension of cellular TPD and for identifying improved agents for therapeutic TPDs.

Our current method, while promising, does have certain limitations that need to be addressed in future iterations. For instance, the *in vitro* unfolding assay exhibits significant variability between biological replicates, which hinders the quantification of results. Enhancements in triggering the reaction within the fluorescent reader could potentially mitigate this issue. Additionally, achieving 100% cellular transfection of P-NADs is currently not feasible and can induce stress in cells. Exploring alternative delivery methods, such as RNA-delivery systems, could offer a solution.

As for all nanobody-based systems, the availability of target-specific nanobodies is less and less of a limiting factor for P-NAD applications, as there are now commercial services for raising them against targets of interest. Moreover, recent advancements in AI-based design of protein binders (Chidyausiku, 2022; Cohen, 2022; Moreno, 2022) indicate that nanobodies could soon be routinely replaced by *in silico* designed modules. Consequently, P-NADs hold the potential to inspire a new class of ubiquitin-independent agents for TPD in both research and therapeutic applications.

## METHODS

### Purifications of proteins

PUX3, PUX4 and P-NADs with N-terminal His tags, CDC48A and p97 with N-terminal GST tags were expressed in *E. coli* BL21 DE3 cells (see **Figure 1D** for sequences). Cells were grown in LB Broth to an OD600 of 0.6-0.8. Protein expression was induced by the addition of 0.5 mM isopropyl b-D-1-thiogalactopyranoside (IPTG) for 16 hr at 18°C. Cells were harvested by centrifugation at 10627 x g for 10 min and resuspended in lysis buffer (50 mM Tris pH 7.5, 500 mM NaCl, 1 mM TCEP). Protease inhibitor, lysozyme, and benzonase were added, and cells were lysed by sonication. Lysates were cleared by ultracentrifugation in a T29-8 x 50 rotor (Thermo) at 68905 x g for 30 min at 4°C.

For CDC48A and p97, supernatants were incubated with Glutathione Sepharose 4B resin for 60 min at 4°C. The resin was washed three times with 10 column volumes of buffer (50 mM Tris pH 8, 500 mM NaCl, 1 mM TCEP, 2 mM EDTA, pH 8). GST tags were removed by incubation with 3C protease overnight at 4C. Proteins were loaded onto a size-exclusion column (Superdex 200) equilibrated in 20mM HEPES, 150mM NaCl, 1mM TECP, pH7.5. Peak fractions were pooled, concentrated, snap-frozen in liquid nitrogen, and stored at -80C. For PUX3, PUX4, and P-NADs, the purification follows a similar protocol. Supernatants were incubated with nickel columns, and eluted by imidazole. Proteins were loaded onto size-exclusion columns for further purification.

### Isothermal Titration Calorimetry (ITC)

The proteins for ITC measurements were dialyzed in the same buffer (20 mM HEPES, 150 mM NaCl, 1 mM TCEP, pH 7.5) overnight. The cell and syringe of ITC (MicroCal PEAQ-ITC and MicroCal ITC 200) were washed with Milli-Q water and then with ITC buffer before protein loading. The instrument was equilibrated to a constant temperature and the cell was stirred continuously to ensure homogeneity. A series of injections were then performed, while the heat change was recorded by the instrument. The ITC measurement started with an initial injection of 0.4 μL, followed by 18 injections of 2 μL each. The experiment was performed at a temperature of 25 °C and a stirring speed of 750 RPM. The spacing between injections was 150 seconds and the injection duration was 6 seconds. The reference power was set to either 5.0 or 10.0 μcal/s. The data were analysed using MicroCal PEAQ-ITC Analysis Software to determine the thermodynamic parameters of the protein-protein interaction, including the dissociation constant (Kd), enthalpy change (ΔH), and stoichiometry (N).

### Fluorescence Unfolding Assay

Assays were carried out at 37°C. A 30 μL reaction contained 200 nM GFP substrate, 366 nM CDC48/p97, 0.5-1.5 mM P-NAD, in unfolding assay buffer: 20 mM HEPES, 150 mM NaCl, 10 mM MgCl2, 5 mM ATP, 1X ATP regeneration buffer (Enzo), pH 7.5. Proteins were mixed and transferred into a black 384-well plate with a clear bottom. The fluorescence intensity was then measured from bottom with excitation at 485 nm and emission at 510 nm by microplate reader (TECAN Infinite® M1000), with an interval of 20 seconds. Relative fluorescence was calculated by normalising the fluorescence signal to that at time 0.

### ATPase Activity Assay

The Malachite Green Phosphate Assay kit (Sigma Aldrich) was used to quantify the production of free phosphate resulting from the hydrolysis of ATP by CDC48 or p97. The assay was carried out at a temperature of 30°C in ATPase buffer (comprising 50 mM HEPES, pH 7.5, 50 mM NaCl, 10 mM MgCl2, 0.5 mM TCEP, and 0.1 mg/mL BSA). Prior to the addition of ATP, Cdc48A or p97 (150 nM), P-NAD4 (500 nM), and GFP (2.5 μM) underwent a pre-incubation of 10 minutes. Subsequently, upon the addition of ATP, the reactions were incubated for 30 minutes. The reaction was halted by introducing the Malachite green reagent and allowing the reaction to incubate for an additional 20 minutes. The absorbance at 640 nm was measured using a Microplate reader (TECAN Infinite M1000 PRO). The amount of phosphate production was determined by the standard curve.

### Plasmid Transfection

Plasmids obtained from Twist Bioscience (up to ∼2000 ng) contained one open reading frame (ORF) under cytomegalovirus (CMV) promoter and selective marker gene of ampicillin resistance. HeLa cells were grown in DMEM medium supplemented with 10% FBS under 5% CO2 and 37ºC temperature conditions. The transfection was performed using Lipofectamine™ 2000 Transfection Reagent (ThermoFisher Scientific) according to the manufacturer’s instructions. In cotransfection experiments, plasmids were mixed in a 1:1 ratio.

### Fluorescence Imaging

Images were obtained with the EVOS M5000 Imaging System, 20x magnification, with the excitation at 470 nm and emission at 525 nm. Images were recorded with High-sensitivity 3.2 MP 2048 x 1536) pixels.

### Western Blot

Transfected cells were harvested on the first day after transfection with a RIPA (Tris buffer, 1 mM PMSF, 1% Triton X). After incubation on ice for 20 minutes a tube with a lysate was centrifuged and a supernatant was analysed further. Gel-electrophoresis was performed in MES buffer and commercially available polyacrylamide gel (ThermoFisher). The transfer of proteins to the PVDF Transfer Membrane (ThermoFisher) was performed with the Pierce™ G2 Fast Blotter. Before usage, the membrane was washed with methanol and Pierce™ 1-Step Transfer Buffer, and the filter layer was washed with Pierce™ 1-Step Transfer Buffer. Staining of bands was done with the SuperSignal West Pico PLUS kit. A signal was collected in the accumulation mode with ChemiDoc XRS+ (Biorad). As a loading control, the signal from anti-GAPDH antibodies was detected.

## ACKNOWLEDGEMENTS

We thank the Bioscience Core Lab and Imaging and Characterization Core Lab at KAUST for their support. For computer time, this research used the resources of the Supercomputing Laboratory at KAUST.

## AUTHOR CONTRIBUTIONS

STA and AS designed and supervised the project. IS supervised the cell experiments. JW, GC, JZ, BH and AS established experimental protocols and collected data. All authors analysed data. STA drafted the manuscript, all authors wrote the Methods section, and commented on the manuscript.

