## Supplementary Figures 1-3 for "P-NADs: *P*UX-based *NA*nobody Degraders for Ubiquitin-Independent Degradation of Target Proteins"

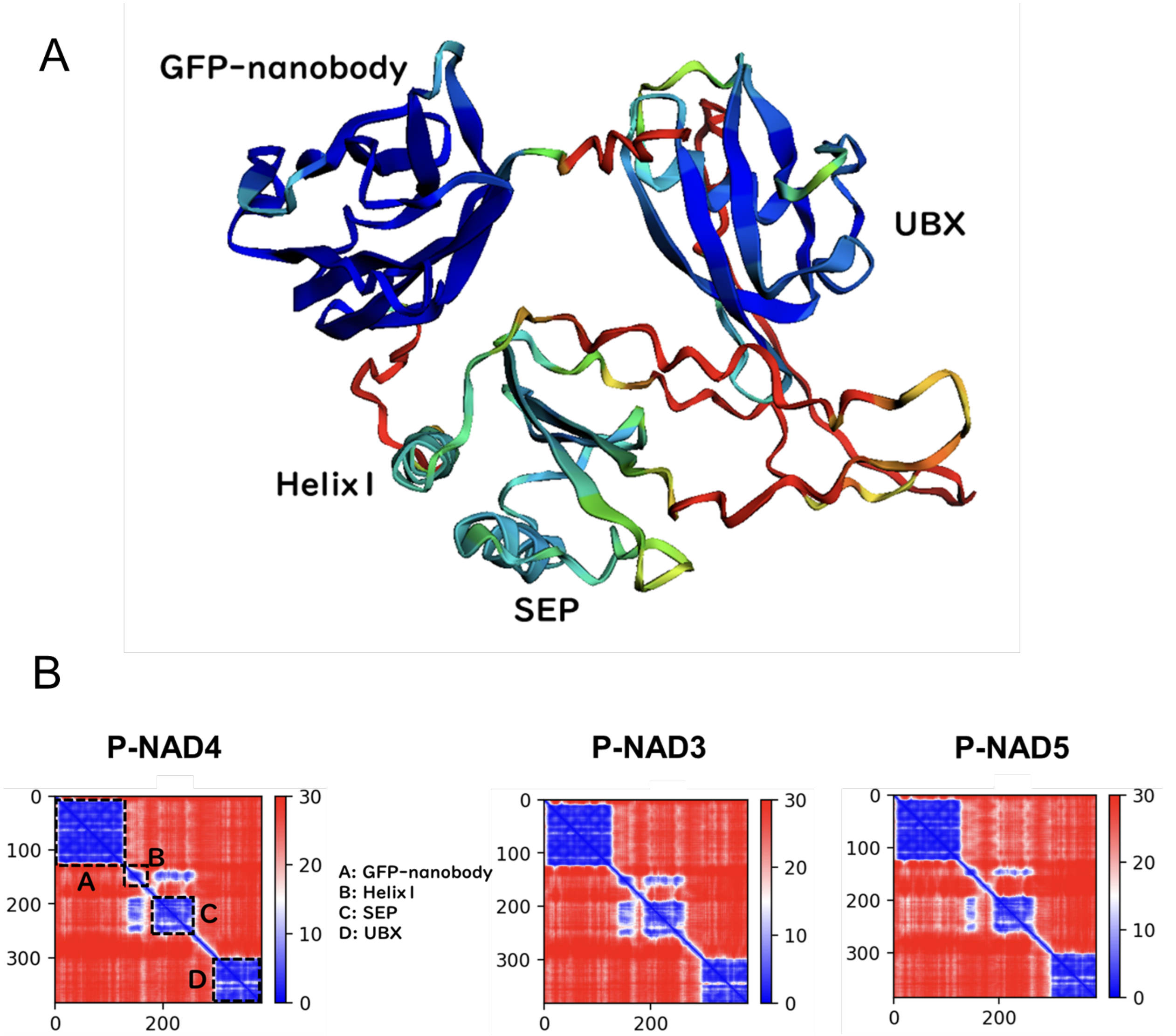


**Supplementary Figure 1. AlphaFold structure prediction of P-NADs.** (**A**) Representative predicted three-dimensional structure of P-NAD4 and predicted aligned error (PAE). (**B**) of P-NAD3, 4, and 5, generated by AlphaFold. The PAE values in Å are shown by a colour gradient from blue (low) to red (high). Domains and motifs are labelled for P-NAD4.

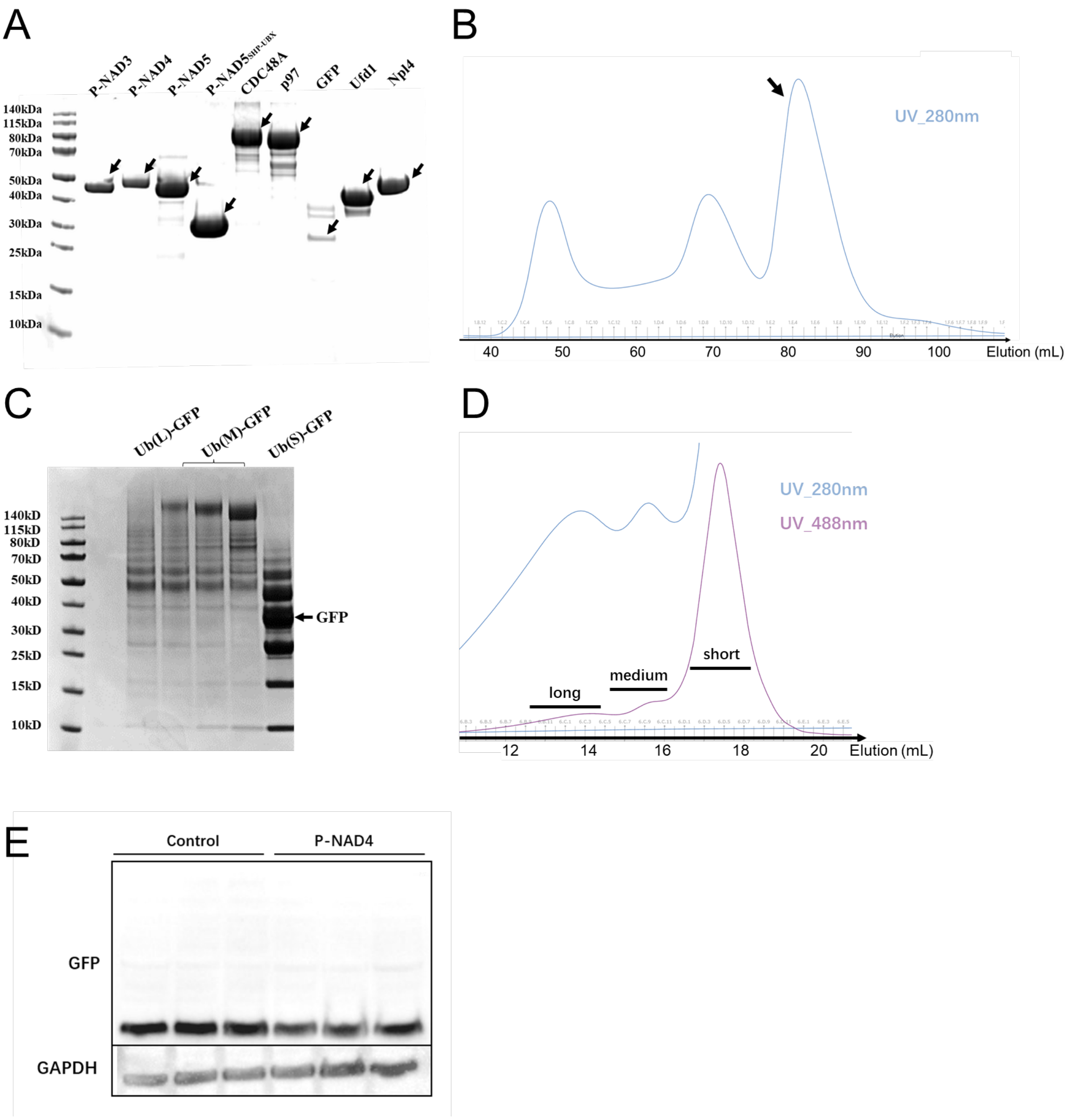


**Supplementary Figure 2. Protein purification and characterization.** (**A**) SDS-PAGE analysis of the purified proteins used in the *in vitro* assays. The target bands are indicated by arrows. The molecular weight markers are identified on the left. (**B**) Size exclusion chromatography (SEC) profile of a representative P-NAD (P-NAD4). The elution volume and the absorbance at 280 nm are shown. (**C**) SDS-PAGE analysis of poly-ubiquitinated GFP (Ub-GFP) after SEC. GFP containing degron sequence was incubated with E1, E2, E3, and ubiquitin, and then separated by SEC. The different lengths of the ubiquitin chains are indicated by L (long), M (medium), and S (short). (**D**) SEC profile of Ub-GFP. The elution volume and the absorbance at 488 nm are shown. The peaks corresponding to the different lengths of the ubiquitin chains are indicated as long, medium or short. (**E**) Uncropped gel from western bolt shown in **Figure 3D** of the total cell lysates, probed with an antibody against GFP. The blot shows the protein level of GFP and the loading control (GAPDH). No higher molecular weight GFP species are apparent, supporting that transfected GFP was not ubiquitinated.

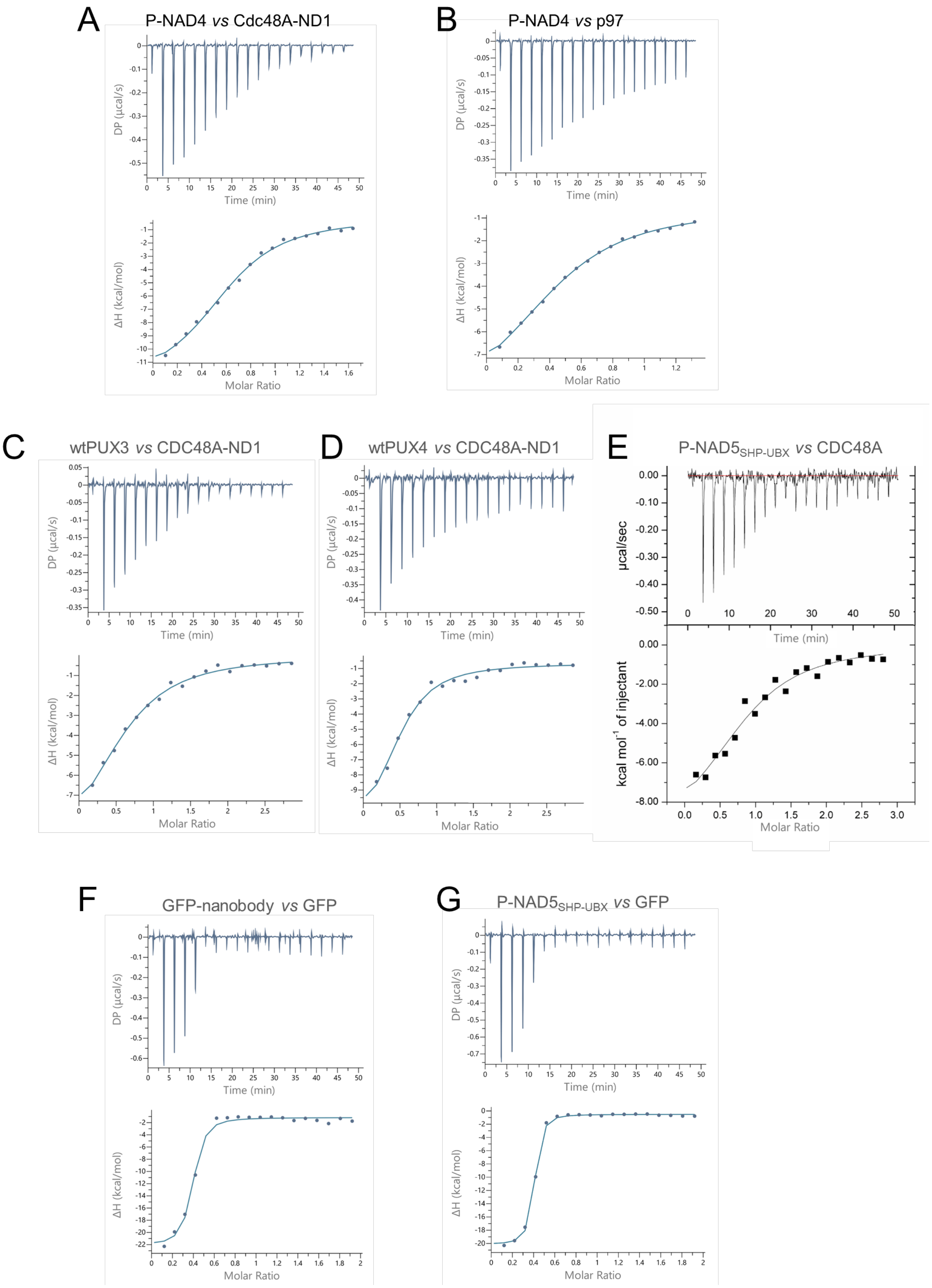
A
P-NAD4 vs Cdc48A-ND1
B
P-NAD4 vs p97
C
wtPUX3 vs CDC48A-ND1
D
wtPUX4 vs CDC48A-ND1
E
P-NAD5SHP-UBX vs CDC48A
F
GFP-nanobody vs GFP
G
P-NAD5SHP-UBX vs GFP
DP (µcal/s)
Time (min)
ΔH (kcal/mol)
Molar Ratio
µcal/sec
kcal mol-1 of injectant

**Supplementary Figure 3. Isothermal titration calorimetry (ITC) results.** (**A**) binding of P-NAD4 vs Cdc48A-ND1; (**B**) P-NAD4 vs p97 (**C**) wtPUX3 vs CDC48A-ND1; (**D**) wtPUX4 vs CDC48A-ND1; (**E**) P-NAD5SHP-UBX vs CDC48A. For each titration, CDC48A or p97 were in the measurement cell, and the degraders were injected. (**F**) GFP-nanobody vs GFP and (**G**) P-NAD5SHP-UBX vs GFP. The thermodynamic parameters, especially the N-value of these titrations are affected by the effect of the GFP chromophore on the absorption coefficient. The upper panels show the raw data of the heat released upon each injection of the P-NADs into the other protein. The lower panels show the integrated and normalised binding isotherms, fitted with a one-site binding model.
